# From concentration to export: resource contrasts and bee traits shape pollinator spillover to crops

**DOI:** 10.64898/2026.08.29.747339

**Authors:** Cristina A. Kita, Isabel Alves-dos-Santos, Michael Hrncir, Renata L. Muylaert, Marco A. R. Mello

**Author notes:** **Corresponding Author:** Cristina A. Kita.

## Abstract

Floral plantings can either concentrate bees or export them to adjacent crops, yet the ecological conditions influencing these outcomes remain unclear. Here, we develop a mathematical model as proof of concept for our previous integrative hypothesis: concentrator and exporter outcomes can arise as alternative, context-dependent outcomes of the same underlying resource-selection process. Using bees as a model and focusing specifically on spillover from floral plantings to crops, we identified resource-specific thresholds separating concentration- and export-favoring conditions. Our model translates differences in relative patch attractiveness into context-dependent concentration and export outcomes and generates resource-specific, testable predictions about the conditions favoring pollinator movement into crops. In our simulations, the concentrator-exporter transition occurred at a lower flowering-intensity contrast than at pollen or nectar contrasts, which suggests that flowering intensity may provide an initial cue for bee movement, whereas nectar and pollen rewards refine or sustain bee responses once crops are perceived as attractive. Spillover thresholds differed among resource contrasts, whereas response steepness varied across bee-trait and community scenarios. Under the model’s trait-sensitivity formulation, predicted spillover probability responded more strongly to flowering contrast for specialists than for generalists; colony size amplified this response, whereas bee richness dampened it. Together, these patterns show how flowering and resource contrasts interact with bee traits and community context to shape predicted spillover. Our results confirm that the concentrator and exporter hypotheses can be understood as context-dependent outcomes of the same ecological process rather than as mutually exclusive alternatives. Experimental tests of the predicted thresholds conducted in the field could reveal when and where floral plantings are most likely to promote bee spillover to crops, potentially supporting crop pollination.

## Introduction

Worldwide, 90% of all flowering plants depend on animal pollination to reproduce (Tong et al. 2023), making the ongoing global pollinator decline a serious threat to ecosystem functioning (Potts et al. 2010, 2016). In addition, through cascading effects on plant diversity and ecosystem structure, the consequences of losing pollinators extend beyond natural ecosystems (Lee et al. 2024).

Many crops also depend on pollinators (Siopa et al. 2024), whose contributions to yield and crop quality translate into higher economic value of agricultural production (Turo et al. 2024). However, intensive agricultural practices are among the major drivers of pollinator decline, as they usually lead to habitat loss and fragmentation (Dicks et al. 2021). This dilemma intensifies the agricultural challenge: how can we increase production while preserving the biodiversity that sustains it? (Sustainable Development Goal 2) (United Nations 2015).

One nature-based solution to the produce vs. conserve dilemma is the adoption of biodiversity-friendly practices, such as the use of floral plantings adjacent to crops, mainly hedgerows and flower strips (Pywell et al. 2015). By providing essential floral resources, floral plantings attract and support pollinators in agricultural landscapes, in addition to boosting ecosystem services, such as the crop pollination service (Nagano et al. 2025). However, they must be carefully managed to ensure this crop pollination goal.

Although floral plantings may facilitate pollinator spillover into the adjacent crop (also known as the “Exporter hypothesis”), they may also concentrate pollinators for themselves (“Concentrator hypothesis”) (Morandin and Kremen 2013). Consistent with both hypotheses, commonly assumed to be mutually exclusive, the effects of floral plantings on crop pollination are highly variable across empirical studies (Lowe et al. 2021). To clarify the mechanisms underlying this apparent inconsistency, and particularly to identify the key factors driving the exporter effect (i.e., pollinator spillover from floral plantings to crops), in a previous study, we proposed a novel integrative hypothesis, based on the principles of a recently developed semantic theory (Mello and Dormann 2025, Mello et al. 2025).

Our integrative hypothesis, termed here the Pollinator Spillover Hypothesis, proposes that the Concentrator and Exporter hypotheses are not mutually exclusive mechanisms but alternative, context-dependent outcomes of the same underlying process: pollinator resource selection between floral plantings and crops (Kita et al. 2026). Within this framework, relative patch attractiveness determines whether predicted movement favors concentration in floral plantings or export to crops, whereas consumer traits determine responsiveness to differences between patches. This formulation connects pollinator movement to resource dissimilarity, which has been proposed as a key driver of consumer–resource interaction structure (Mello et al. 2025).

Formalizing the Concentrator and Exporter hypotheses as context-dependent outcomes of a common resource-selection mechanism provides a unified basis for identifying the conditions that shift bee movement from concentration in floral plantings toward export to crops. Examining how these transition thresholds vary with resource contrasts, consumer traits, and ecological context provides a mechanistic basis for predicting when concentration or export should emerge, generating empirically testable expectations about when floral plantings are most likely to promote pollinator movement into crops. Once empirically evaluated, these predictions could inform the design and management of floral plantings that support both pollinator conservation and crop pollination.

In this context, given that bees are the main pollinators of most crops (Requier et al. 2023), our objective was to develop a theoretical proof of concept for the Pollinator Spillover Hypothesis and derive empirically testable predictions that could ultimately guide the sustainable management of floral plantings to support pollinator conservation while enhancing their potential contribution to crop pollination. Using a computational modeling approach, we: (1) formalized pollinator concentration and export as alternative outcomes of a common resource-selection mechanism; (2) evaluated how ecological and environmental variables translated this mechanism into variation in predicted spillover; and (3) estimated resource-specific movement thresholds under different bee-trait and community contexts.

## Methods

### Model semantics

We developed a computational model as a proof of concept (Servedio et al. 2014) for the Pollinator Spillover Hypothesis by formalizing its logical structure, evaluating the consequences of its component assumptions, and generating testable predictions in silico. We used a mathematical modeling approach (Shoemaker et al. 2021) to determine how resource contrasts and bee traits translate a common resource-selection mechanism into predicted variation in bee spillover from floral plantings to crops. The model was developed in conceptual connection with the Integrative Theory of Interaction Networks, which identifies resource dissimilarity and consumer specialization as key drivers of consumer–resource interaction structure (Mello et al. 2025). We apply these principles to spatial resource selection by examining how resource contrasts and consumer traits shape movement between adjacent patches.

Because our objective was not to reproduce the full temporal dynamics of bee populations, but rather to identify the mechanisms and transition points underlying bee spillover from floral plantings to crops, we intentionally simplified demographic processes. Bee communities were represented as behavioral snapshots in which individuals redistribute between habitats according to ecological and environmental conditions, rather than through explicit processes of population growth, mortality, and migration. This approach allowed us to focus on the emergence of spillover and transition behavior while maintaining a manageable level of complexity. Temporal variation was incorporated indirectly through phenological snapshots representing different stages of flowering season rather than through explicit population dynamics. Before constructing the model, we defined a modeling workspace combining empirical data and simulated values within ecologically plausible ranges (online repository: Table S1).

### Modeling workspace

To identify general ecological patterns rather than reproduce in detail any particular agricultural system, we simulated bee movement across a broad range of landscape conditions and climatic settings, designed to represent variation among agricultural environments. We simulated 10,000 individual bees across 500 landscapes in different climatic zones, with 20 bees per landscape. Each landscape was assigned to a tropical, subtropical, or temperate climatic zone, defined by scaled temperature and precipitation ranges, and included a floral planting, a crop type, natural habitat fragments, and a bee community (Figure 1). In our simulation, bee abundance was held constant so that variation in spillover dynamics reflected environmental and behavioral factors rather than differences in population size.

**Figure 1.**
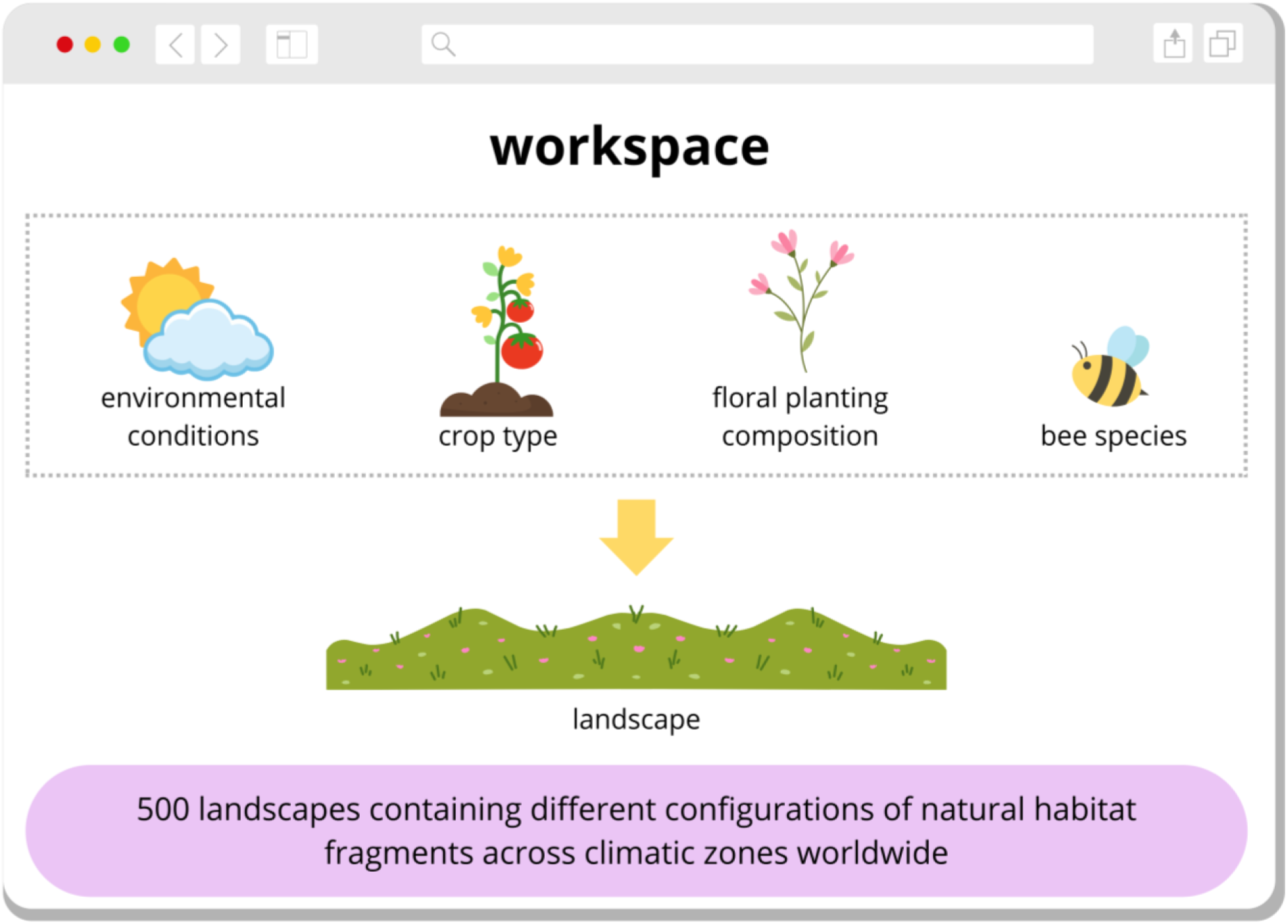
Reproducible computational modeling framework. A set of ecological factors was used to simulate bee movement across 500 landscapes varying in natural habitat configuration and climatic conditions worldwide. Illustration created by the authors using elements licensed under the Canva Content License Agreement.

Floral plantings and crops varied in nectar and pollen production, with values simulated from empirical data and local stochastic deviations to reflect ecological realism (sensu Levins 1966). Natural habitat fragments also varied in number, plant species richness, and distance from floral plantings, further contributing to landscape heterogeneity (online repository: Table S1).

Building on the simulated spatial structure, we incorporated plant phenological variation by representing each simulation as a snapshot of early, mid, or late flowering periods. These phases resulted in different flowering intensities for crops and floral plantings, which in turn influenced nectar and pollen availability, patch attractiveness, and bee movement probability. Full phenological details are provided in Appendix S1.

### Main factors

We focused on the four key factors that seem to influence bee movement from floral plantings to crops: bee species, floral planting composition, crop type, and environmental conditions (see Appendix S1: Figure S1). For each factor, we defined key traits following ecological guiding principles and translated them into operational variables used directly in the model equations (Figure 2). Below, we briefly summarize the traits included in the model, while full parameterization details are provided in the online supplement (Appendix S1).

**Figure 2.**
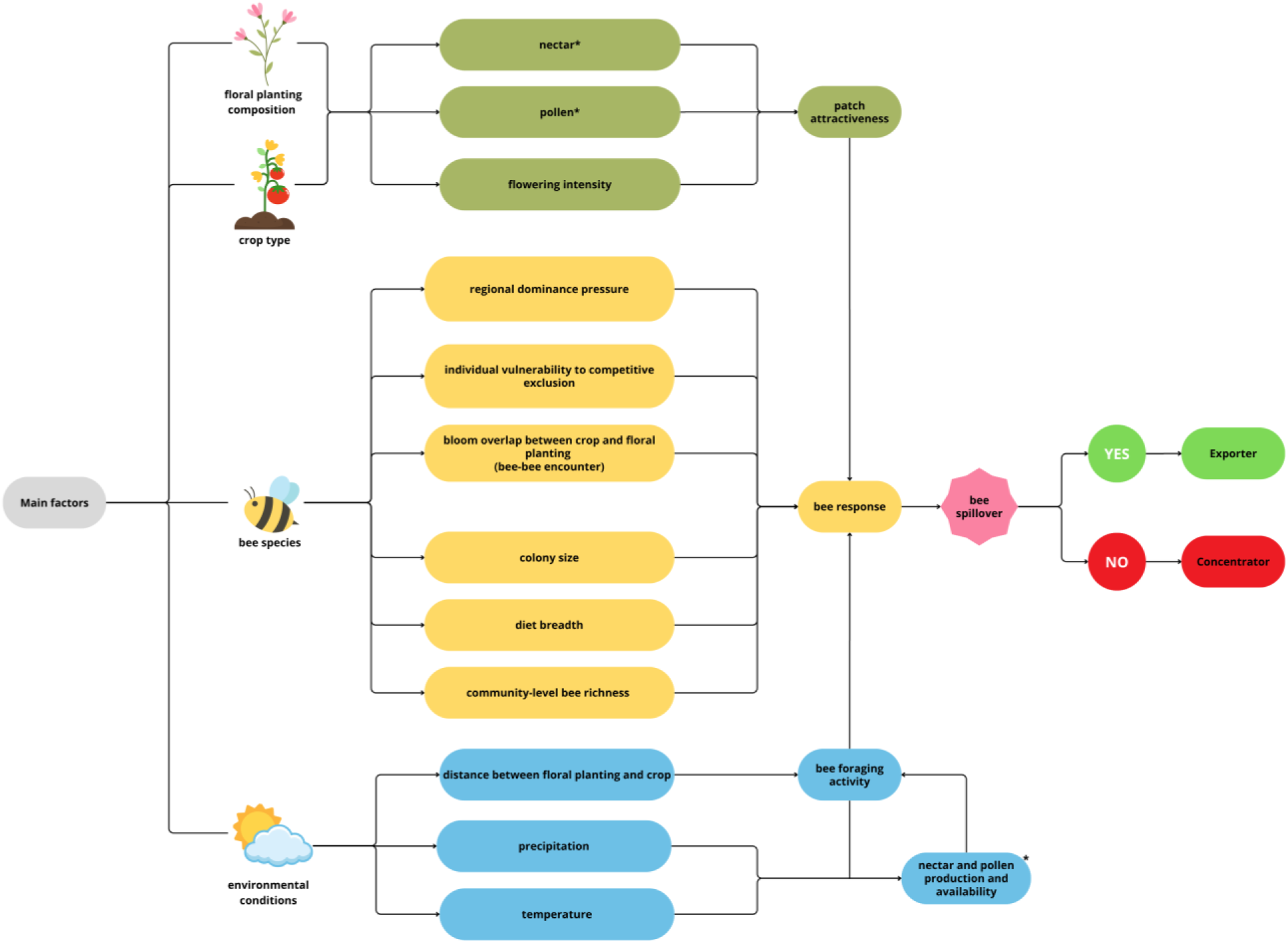
Conceptual framework of the probabilistic model of bee spillover from floral plantings to crops. The model articulates operational variables embedded within the key factors that influence bee spillover towards crops, as well as their relationships: floral planting composition, crop type, bee species, and environmental conditions. Illustration created by the authors using elements licensed under the Canva Content License Agreement.

#### Factor 1: floral planting composition

Floral plantings varied in plant richness, flowering dynamics, resource availability, and resource accessibility. Plant richness represented variation in the number of plant species. Differences among species in flowering duration were represented at the community level: floral plantings containing a greater proportion of long flowering species had a broader flowering window and therefore remained in bloom over a larger portion of the flowering period. The effective availability of resources (nectar and pollen) is represented by the relative number of open flowers at a given time (flowering intensity) within the flowering window. Resource availability was also modified by climatic conditions, including temperature and precipitation. Resource accessibility broadly represented constraints imposed by factors such as floral morphology and rainfall. However, we did not explicitly simulate particular flower morphologies or rainfall regimes, bee–flower trait matching, or specialized foraging strategies such as nectar robbing (Irwin et al. 2010).

#### Factor 2: crop type

Crops were simulated using the same resource-based framework as floral plantings, but were represented as monocultures, with plant richness fixed at one species. As in floral plantings, crop attractiveness depended on flowering intensity, nectar and pollen availability, and resource accessibility, with resource availability and accessibility modified by climatic conditions.

#### Factor 3: bee species

Bee species were characterized by foraging range, competitive dominance, colony size, and diet breadth. Foraging range captured broad interspecific variation in movement capacity, which is commonly associated with body size and sociality (Greenleaf et al. 2007, Grüter and Hayes 2022), whereas competitive dominance represented the ability to monopolize floral resources or displace competitors from a resource source (Lichtenberg et al. 2009). Colony size and diet breadth jointly determined sensitivity to differences in patch attractiveness. In the trait-assignment step, sociality was inferred from colony size, and diet breadth was assigned conditionally on sociality, with solitary bees more likely to have narrower diet breadths (specialists) and social or eusocial bees more likely to have broader diet breadths (generalists). In the sensitivity function, diet breadth determined baseline sensitivity to resource contrasts, whereas colony size acted as a proxy for colony-level resource demand and recruitment-mediated amplification in social bees. Thus, solitary bees, whose colony size was set to one, had no colony-level amplification, while bees with narrower diet breadths had higher baseline sensitivity to resource contrasts.

#### Factor 4: environmental conditions

Environmental context included climate and landscape structure. Temperature and precipitation varied among tropical, subtropical, and temperate zones and modified floral resources, accessibility, and bee foraging conditions. The diversity and connectivity of natural fragments, together with crop distance, represented spatial context, while climate and floral planting diversity contributed to variation in bee richness. These factors influenced spillover through both movement feasibility and changes in resources and bee communities. In addition to these measured environmental and community differences, each simulated region received a latent random intercept drawn from a normal distribution with mean 0 and standard deviation 0.4 on the logit scale. This term represented unmeasured differences among landscapes, such as variation in pesticide exposure, habitat permeability, or other regional characteristics not explicitly included in the model. All observations from the same region shared the same intercept. Because this term was generated independently of climatic zone, it increased heterogeneity among landscapes within each zone without removing the broad climatic differences among tropical, subtropical, and temperate regions.

### Core equation

After simulating the variables of each main factor, we built a bee spillover probability equation that was modeled as a function of the relative attractiveness of the crop and the floral planting, incorporating resource availability, resource accessibility, competitive dominance, bee behavioral sensitivity to patch resource contrast, and the spatial constraint imposed by the distance between the floral planting and the crop.

To estimate bee movement, we modeled crop and floral planting attractiveness separately, using nectar and pollen as the core resource components because they are broadly used as energetic and nutritional rewards. For each patch, effective resource availability depended on flowering intensity, raw nectar and pollen supply, climatic conditions, and floral accessibility. Accessibility reflected how easily bees could reach floral rewards, varying with flower morphology and declining under rain.

Accessible nectar and pollen were combined into a resource index used to estimate baseline attractiveness for each patch. Crop attractiveness was then reduced under high competitive dominance, especially when crop and floral planting bloom overlapped, dominant bees were common, and the focal bee was vulnerable to displacement. Thus, patch attractiveness reflected both resource availability and access.

Bee responses to differences in attractiveness between crops and floral plantings were scaled by a trait-based sensitivity parameter. This sensitivity increased with dietary specialization and colony size. Sensitivity was then adjusted by bee richness and regional climatic conditions, allowing community context and climate to modulate how sharply bees responded to differences in patch attractiveness.

Finally, bee movement was modeled as a probabilistic choice between floral plantings and crops based on their relative attractiveness. If the crop was beyond the bee’s flight range, spillover probability was set to zero. Otherwise, the probability of spillover increased when crop attractiveness exceeded floral planting attractiveness, with the strength of this response depending on bee sensitivity. Thus, the resource-selection component of spillover probability is:

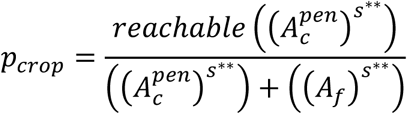

where *p_crop_* is the probability of bee spillover from floral plantings to crops; 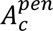 is crop attractiveness after accounting for nectar and pollen availability, floral accessibility, climate, and competitive dominance; *A_f_* is floral planting attractiveness after accounting for nectar and pollen availability, floral accessibility, and climate; *s*^∗∗^ is the adjusted bee sensitivity parameter, which determines how strongly bees respond to differences in attractiveness between patches; and *reac*ℎ*able* indicates whether the crop is within the bee’s flight range. In this formulation, the numerator represents the attractive force of the crop, whereas the denominator represents the combined attractiveness of the crop and floral planting. Intermediate equations and implementation details are provided in Appendix S1.

This equation describes the resource-selection component of bee movement. To account for unmeasured ecological differences among landscapes, such as pesticide exposure, local disturbance, and the availability of alternative floral resources not represented in the model, a region-specific random intercept was added to the logit of the spillover probability for bees able to reach the crop. This allowed regions to differ in their baseline tendency toward export without changing the relative-attractiveness mechanism described above. For bees unable to reach the crop, spillover probability remained zero. Further details are provided in Appendix S1.

We represented trait- and context-mediated resource selection by allowing diet breadth, colony size, and bee richness to modify bee sensitivity to contrasts in patch attractiveness, with the direction of these relationships specified a priori based on ecological expectations. Movement probability was then determined by relative patch attractiveness, integrating flowering intensity, nectar and pollen availability, resource accessibility, competitive effects, spatial reachability, and trait-mediated sensitivity across heterogeneous phenological, climatic, and landscape conditions. This formulation allowed us to quantify how resource contrasts and ecological context affected predicted bee movement from floral plantings to crops.

To assist in optimizing and debugging the code used to implement our mathematical model, we used generative AI tools, specifically ChatGPT (OpenAI, GPT-5.5; https://chat.openai.com/), Codex (OpenAI, GPT-5.5; https://openai.com/codex/), and Claude (Anthropic, Claude Opus 4.6; https://claude.ai/). All AI-assisted outputs were reviewed, tested, and validated by the authors, who take full responsibility for the final code, analyses, and conclusions.

### Model checking and characterization of the movement response

Before conducting the ecological analyses, we performed model checks to verify that the simulations reproduced the behavior specified by the movement equation. We confirmed that resource indices remained within their intended ranges, that crop-floral planting attractiveness contrasts included both negative and positive values, and that these contrasts occurred across flowering-period phases. For reachable crops, we also evaluated the calibration of the simulated binary outcomes against the probabilities generated by the movement equation.

The simulations included crop locations both within and beyond the bees’ foraging ranges. When the distance exceeded a bee’s foraging range, the crop was classified as unreachable, and its spillover probability was set to zero. We then restricted the statistical analysis to cases in which the crop was reachable, so the binary outcome represented behavioral movement rather than spatial impossibility (i.e., spillover probability = 0): 0 for remaining in the floral planting and 1 for moving to the crop.

As an implementation check, we compared logit, probit, complementary log-log, and cauchit links using cross-validation and calibration metrics. Logit and probit showed similar performance, and we retained the logit link because it was consistent with the mathematical formulation of the movement model. Details of model checking and calibration are provided in Appendix S1.

### Statistical modeling and sensitivity analysis

We used a binomial generalized linear mixed model (GLMM) with a logit link to characterize how the components of the movement model contributed to variation in simulated bee spillover. The GLMM quantified the direction, magnitude, and context dependence of these contributions across the simulated parameter space. We restricted the analysis to crops within the bees’ foraging ranges and grouped predictors according to the model structure (Appendix S1: Figure S2). We then compared changes in deviance among nested models to estimate the relative contribution of each model component (Nelder and Wedderburn 1972, Bolker et al. 2009). Details of fixed and random variables and sensitivity analysis are provided in Appendix S1.

### Threshold analysis

Based on the sensitivity analysis and GLMM estimates, we selected flowering, nectar, and pollen contrasts as focal predictors of spillover. For each contrast, we asked two complementary questions: (1) at what value does movement to the crop become more likely than concentration in the floral planting, and (2) how rapidly does this probability change around that value?

We addressed the first question by estimating the threshold position. We defined the spillover threshold as the value of a resource contrast at which the predicted probability of movement to the crop reached 0.5. Values below the threshold favored concentration in the floral planting, whereas values above it favored export to the crop. This threshold represents a decision boundary and does not imply an abrupt or discontinuous change in movement probability. For bees able to reach the crop, a probability of 0.5 corresponds approximately to equal overall adjusted attractiveness at the population-average level, because the regional random intercept has a mean of zero. Within a particular region, however, latent regional heterogeneity may shift this decision boundary. However, a threshold expressed for flowering, nectar, or pollen does not indicate that the two patches are equal in that individual resource. Instead, it indicates how large that particular resource contrast must be for the two patches to become equally attractive after accounting for all other conditions represented in the model.

We addressed the second question by estimating response steepness, which describes how rapidly spillover probability changes as a resource contrast crosses its threshold. A steep response indicates a rapid shift from concentration-to export-favoring conditions, whereas a gradual response indicates a smoother change in movement probability. We estimated marginal thresholds to describe the average system-level threshold for each resource contrast and conditional response slopes to evaluate how response steepness varied among colony-size, bee-richness, and diet-breadth scenarios. Thresholds are reported in standardized units, and their uncertainty was quantified using a parametric bootstrap with 1,000 iterations. Full estimation details are provided in Appendix S1.

## Results

### Movement response to relative patch attractiveness

Model checks showed that the simulated movement outcomes followed the resource-selection rule specified in the model. For crops within the bees’ flight range, spillover probability increased as crop attractiveness became greater than floral-planting attractiveness. When adjusted crop and floral-planting attractiveness were similar, the bee-weighted resource contrast approached zero and, on average across regions, movement toward either patch was approximately equally likely. As the contrast became more negative, concentration in floral plantings became more likely; as it became more positive, export to crops became more likely (Figure 3). This result confirms the internal consistency of the model implementation and provides the reference point for estimating how floral resources and bee traits shape spillover thresholds.

**Figure 3.**
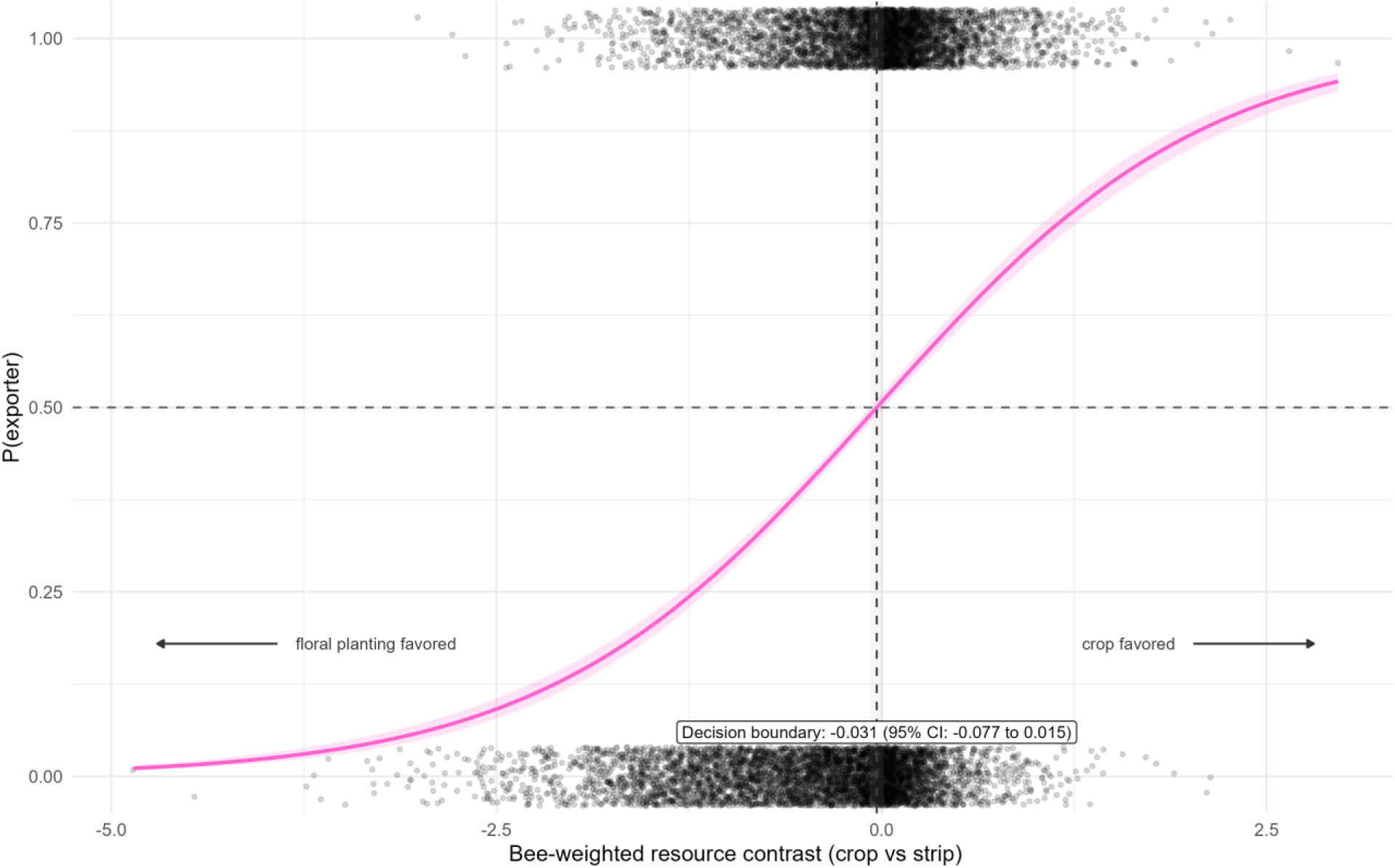
Internal consistency of the simulated movement response. Relationship between bee-weighted resource contrast and the probability of bee spillover to the crop. Bee-weighted resource contrast was defined as adjusted bee sensitivity multiplied by the log ratio of crop to floral-planting attractiveness. Negative values favor concentration in floral plantings, whereas positive values favor movement to crops. Points represent simulated binary outcomes, the solid curve represents the fitted logistic response, and the shaded area represents its 95% confidence interval. The vertical dashed line indicates the fitted decision boundary (x = −0.031; 95% CI: −0.077 to 0.015), which was consistent with the theoretical boundary at x = 0, where the two patches have equal adjusted attractiveness and P(export) = 0.5. This figure evaluates the internal consistency of the model implementation rather than independently testing the logistic functional form.

### Resource contrasts and bee traits shape spillover responses

Within the simulated system, trait-mediated differences in responses to resource contrasts were concentrated primarily in flowering contrast. For the reference diet-breadth group, nectar and pollen contrasts showed no clear effects (nectar: β = 0.04, p = 0.60; pollen: β = 0.12, p = 0.16), whereas flowering contrast had a small positive effect (β = 0.17, p = 0.04). Mesolectic (β = −0.19, p = 0.04) and oligolectic bees (β = −0.25, p = 0.01) had lower baseline spillover than superpolylectic bees. Within oligolectic bees, conditional responses were positive for nectar (β = 0.18, p < 0.001), pollen (β = 0.19, p < 0.001), and flowering contrasts (β = 0.75, p < 0.001). However, compared with superpolylectic bees, the increase in response was statistically clear only for flowering contrast (interaction β = 0.58, p < 0.001), whereas interactions with nectar and pollen showed no clear differences among these diet-breadth groups. Colony size amplified the response to flowering contrast (β = 0.11, p < 0.001), while its interaction with pollen was marginal (β = 0.04, p = 0.070) and its interaction with nectar was not supported. Bee richness dampened the response to flowering contrast (β = −0.07, p = 0.001) but showed no clear interactions with nectar or pollen. Flowering overlap reduced spillover (β = −0.13, p < 0.001), whereas competitive traits, temperature, precipitation, and distance relative to flight range showed no clear direct effects. Resources and phenology, together with their interactions with bee-sensitivity traits, dominated the block-based sensitivity analysis, accounting for 83.9% of the relative contribution to model fit across block-removal tests (Appendix S1: Figure S3). Bee-sensitivity traits and their interactions with resource contrasts accounted for a further 12.1%. Thus, predicted spillover was structured primarily by resource contrasts and by how bee traits and community context modified responses to those contrasts.

### Threshold analysis

The sensitivity analysis identified resources and phenology, together with their interactions with bee-sensitivity traits, as the most influential component of model fit. We therefore estimated marginal thresholds for flowering, pollen, and nectar contrasts and evaluated conditional response slopes across diet-breadth, colony-size, and bee-richness scenarios.

Marginal threshold positions differed among resource contrasts. Predicted spillover probability crossed 0.5 at 0.41 standard deviations for flowering contrast (95% CI: 0.31–0.52), 1.50 for pollen contrast (95% CI: 1.03–2.30), and 1.85 for nectar contrast (95% CI: 1.24–2.98). Thus, flowering contrast produced the lowest threshold, whereas pollen and nectar required larger crop-floral planting contrasts.

Conditional analyses showed that response steepness to flowering contrast increased with dietary specialization. Spillover odds increased by 19% in superpolylectic bees (OR = 1.19; 95% CI: 1.01–1.39; p = 0.038), by 49% in polylectic bees (OR = 1.49; 95% CI: 1.37–1.62; p < 0.001), by 75% in mesolectic bees (OR = 1.75; 95% CI: 1.61–1.90; p < 0.001), and by 113% in oligolectic bees (OR = 2.13; 95% CI: 1.95–2.31; p < 0.001). Nectar and pollen contrasts showed similar but weaker conditional patterns, although differences in their slopes among diet-breadth groups were not clearly supported. Colony size further amplified response steepness to flowering contrast, whereas increasing bee richness dampened it. Because diet breadth, sociality, and colony size covary in natural bee communities, these conditional patterns provide testable predictions about their joint influence on crop-directed movement. Full conditional estimates are provided in Appendix S1.

Together, these results reinforce that bee spillover was not driven simply by higher resource availability in crops. Instead, predicted movement reflected resource and phenological contrasts between crops and floral plantings, particularly flowering contrast, together with variation in how bee traits and community context shaped responses to those contrasts. Thus, crops became more likely to attract bees not only by offering relatively stronger floral cues and rewards, but also because bee species differed in their sensitivity to those differences.

## Discussion

Our mathematical model provides a theoretical proof of concept for the Pollinator Spillover Hypothesis, showing that pollinator concentration in floral plantings and export to crops can be understood as context-dependent outcomes of a shared resource-selection mechanism. This reframing shifts the central question from whether floral plantings concentrate or export pollinators to when each outcome should emerge. The simulations identified distinct thresholds for flowering, pollen, and nectar contrasts, with flowering producing the lowest threshold. Responses to flowering contrast were further modified by diet breadth, colony size, and bee richness. Accordingly, spillover depended not only on resource differences between patches, but also on which bees encountered those differences and within what community context. These predictions provide a mechanistic basis for empirical tests of when floral plantings are most likely to promote crop visitation. In the following sections, we discuss how these predictions can be empirically tested and how they could guide the sustainable management of floral plantings to support pollinator conservation while enhancing their potential contribution to crop pollination.

Within the simulated parameter space, many simulated observations occurred near a resource contrast of zero (Figure 3), where crops and floral plantings had similar adjusted attractiveness and neither concentration nor export was strongly favored. This pattern suggests a temporal management hypothesis: promoting crop-directed spillover may therefore depend on creating an appropriate temporal contrast between patches. In other words, floral plantings could sustain bee communities before and after crop flowering, while crops become relatively more attractive during the period when pollination is required. This does not imply removing floral resources during crop bloom. Rather, the amount, composition, accessibility, and flowering phenology of floral plantings could be selected to complement crop resources without strongly favoring bee concentration away from crops. Because responses to flowering contrast varied particularly with diet breadth, colony size, and bee richness, effective management would also need to account for the resource requirements and foraging responses of the local bee community. The resource-specific thresholds identified by our model provide quantitative predictions that can be tested when designing such temporal arrangements.

The lower threshold for flowering contrast suggests that relatively small differences in flowering intensity can initiate changes in bee movement, whereas pollen and nectar require larger contrasts. Flowering may therefore provide an initial visual and olfactory cue attracting bees to crops, as observed in oligolectic bees (Burger et al. 2010), honeybees (Rachersberger et al. 2019), and bumblebees (Sommer et al. 2022), while sufficient nectar and pollen rewards help sustain crop visitation. Whether these contrasts produce spillover nevertheless depends on how individual species perceive and respond to resources and how these responses scale to the community level (Bauer et al. 2017). Consistent with this interpretation, exporter effects have been reported in abundantly flowering crops, including almond, apple, sunflower, field mustard, and oilseed rape (Kita et al. 2026), which may combine strong floral signals with sufficient rewards to sustain crop-directed bee movement.

More broadly, our results shift the focus from what is planted to how floral plantings function within their local ecological context. Their effectiveness may depend less on a particular crop or a fixed floral planting composition than on whether management creates the conditions needed to sustain bee populations and promote their movement into crops when pollination is required. This context-specific management should consider both resource complementarity and temporal continuity.

Management should also consider the resource requirements of the local bee community, including whether bees obtain primarily nectar, pollen, or both from the crop. Seasonal constraints of each context should likewise guide plant selection in floral plantings. In addition, because floral plantings are used by organisms other than pollinators, managers should assess which pest species they could attract or support and whether these pests could subsequently spillover into the crop (Rodríguez-Gasol et al. 2026). Management strategies should then seek to minimize these potential disservices without compromising the resources provided to pollinators.

Another important consideration is to survey the local bee community before designing or managing floral plantings. Our results indicate that responses to resource differences between patches, particularly differences in flowering intensity, depend on diet breadth and community richness. For social and eusocial bees, colony size can further modify responses to flowering contrast by representing colony-level resource demand. In the simulations, diet breadth was assigned conditionally to sociality, with solitary bees more likely to be specialists and social bees more likely to be generalists. Therefore, identifying the species present in the landscape can help infer their ecological traits and guide predictions about how they will explore the resources offered by floral plantings and crops. Resource availability alone may therefore be insufficient to predict spillover. Floral planting management should consequently be informed by local bee surveys and account for both the resources available and the ecological traits and foraging behaviors of the species expected to provide crop pollination.

Although we focus on bee movement towards crops, pollinator movement should not be viewed as a unidirectional process. Rather, pollinators may move among native habitats, floral plantings, and crops, linking these habitats through foraging activities (Kratschmer et al. 2026). If facilitated by appropriate floral planting management, movement from floral plantings to crops could therefore be accompanied by return movement from crop areas or floral plantings back to natural habitats, creating positive feedback that benefits both agricultural production and ecosystem functioning.

It is important to note that bridging theory and practice will require testing those thresholds under real-world conditions. Our estimated thresholds provide a general guide for identifying the conditions that may promote bee spillover, rather than universal management prescriptions. Translating them into effective recommendations will require empirical validation in specific crops through experimentation, accounting for their floral resources, pollinator communities, and local environmental conditions. Crop-specific thresholds could then provide a more reliable basis for managing floral plantings to support both pollinator conservation and crop production.

Threshold-based approaches can also provide a useful framework for anticipating how pollination dynamics may change under future scenarios. Because thresholds predicted bee movement changes from favoring concentration in floral plantings to favoring export to crops, the general structure of our mathematical model makes it adaptable to specific ecological contexts and suitable for investigating threshold conditions that determine when floral plantings promote or reduce crop-directed spillover. In this way, the model could help explore how changes in community composition, resource availability, or environmental conditions alter the likelihood of bee movement into crops. For example, since rising temperatures affect plant-pollinator interactions (Freimuth et al. 2022, Peng et al. 2025), investigating how floral planting-bee-crop dynamics respond to environmental change, and the consequences for crop pollination, is a promising approach.

## Conclusion

The Concentrator and Exporter hypotheses can be represented not as mutually exclusive mechanisms but as alternative context-dependent outcomes of a common resource-selection process. Our model predicts distinct resource-specific transition points separating concentration-and export-favoring conditions, while bee traits and ecological context modify the strength and sharpness of these responses. Flowering, pollen, and nectar contrasts therefore provide a mechanistic basis for predicting spillover across context-specific scenarios. Testing these predictions under real-world conditions can guide the design of floral plantings that promote crop-directed bee movement while supporting pollinator conservation.

## Acknowledgments

We are deeply grateful to Tereza Cristina Giannini, Paula Prist, and Silvana Buzato for their insightful suggestions during CAK’s Qualifying Exam. Astrid Kleinert’s invaluable insight and advice helped us see the bigger picture and put CAK’s Ph.D. project in perspective. Generative AI tools were used to assist with code optimization and debugging, as described in the Methods. All AI-assisted outputs were reviewed and validated by the authors, who take full responsibility for the work.

## Funding

CAK thanks the Coordination for the Improvement of Higher Education Personnel (CAPES, 8888.802356/2023-00), Graduate School in Ecology of the University of São Paulo (PPGE/IB-USP), and São Paulo Research Foundation (FAPESP, 2023/17728-9) for the Ph.D. scholarships. RLM is supported by an Australian Research Council Australian Laureate Fellowship (FL240100037) funded by the Australian Government. MARM was supported by grants, fellowships, and scholarships given to him and his team by the Alexander von Humboldt Foundation (AvH, 1134644), São Paulo Research Foundation (FAPESP, 2023/03083-6, 2023/02881-6, and 2023/17728-9), National Council for Scientific and Technological Development (CNPq, 305204/2024-6), and Consulate General of France in São Paulo. IAS is supported by the National Council for Scientific and Technological Development (CNPq, 310659/2026-4) grant.

## Conflict of Interest Statement

The authors declare no conflicts of interest.

## Notes

### Competing Interest Statement

The authors have declared no competing interest.

## References

Bauer, A. A., M. K. Clayton, and J. Brunet. 2017. Floral traits influencing plant attractiveness to three bee species: Consequences for plant reproductive success. American Journal of Botany 104:772–781.

Bolker, B. M., M. E. Brooks, C. J. Clark, S. W. Geange, J. R. Poulsen, M. H. H. Stevens, and J. S. S. White. 2009. Generalized linear mixed models: a practical guide for ecology and evolution. Trends in Ecology and Evolution 24:127–135.

Burger, H., S. Dötterl, and M. Ayasse. 2010. Host-plant finding and recognition by visual and olfactory floral cues in an oligolectic bee. Functional Ecology 24:1234–1240.

Dicks, L. V., T. D. Breeze, H. T. Ngo, D. Senapathi, J. An, M. A. Aizen, P. Basu, D. Buchori, L. Galetto, L. A. Garibaldi, B. Gemmill-Herren, B. G. Howlett, V. L. Imperatriz-Fonseca, S. D. Johnson, A. Kovács-Hostyánszki, Y. J. Kwon, H. M. G. Lattorff, T. Lungharwo, C. L. Seymour, A. J. Vanbergen, and S. G. Potts. 2021. A global-scale expert assessment of drivers and risks associated with pollinator decline. Nature Ecology and Evolution 5:1453– 1461.

Freimuth, J., O. Bossdorf, J. F. Scheepens, and F. M. Willems. 2022. Climate warming changes synchrony of plants and pollinators. Proceedings of the Royal Society B: Biological Sciences 289.

Greenleaf, S. S., N. M. Williams, R. Winfree, and C. Kremen. 2007. Bee foraging ranges and their relationship to body size. Oecologia 153:589–596.

Grüter, C., and L. Hayes. 2022. Sociality is a key driver of foraging ranges in bees. Current Biology 32:5390–5397.e3.

Irwin, R. E., J. L. Bronstein, J. S. Manson, and L. Richardson. 2010. Nectar robbing: Ecological and evolutionary perspectives. Annual Review of Ecology, Evolution, and Systematics 41:271–292.

Kita, C. A., I. Alves-dos-Santos, M. Hrncir, and M. A. R. Mello. 2026. From conservation to crop pollination: an ecological synthesis of floral planting effects on bee activity within crops. Biological Conservation 317:111807.

Kratschmer, S., E. Ockermüller, V. S. Scharnhorst, J. Neumayer, K. Pascher, C. Hainz-Renezeder, N. Sauberer, T. Frank, and B. Pachinger. 2026. Bee – Plant networks in agricultural landscapes are enhanced by increased landscape diversity and agri-environmental measures. Agriculture, Ecosystems & Environment 399:110163.

Lee, Y. D., T. Yokoi, and T. Nakazawa. 2024. A pollinator crisis can decrease plant abundance despite pollinators being herbivores at the larval stage. Scientific Reports 14:18523.

Levins, R. 1966. The strategy of model building in population biology. American Scientist 54:421–431.

Lichtenberg, E. M., V. L. Imperatriz-Fonseca, and J. C. Nieh. 2010. Behavioral suites mediate group-level foraging dynamics in communities of tropical stingless bees. Insectes Sociaux 57:105–113.

Lowe, E. B., R. Groves, and C. Gratton. 2021. Impacts of field-edge flower plantings on pollinator conservation and ecosystem service delivery – A meta-analysis. Agriculture, Ecosystems & Environment 310:107290.

Mello, M. A. R., and C. F. Dormann. 2025. We need to talk about the topology of interaction networks. Oecologia Australis 29:190–195.

Mello, M. A. R., S. E. Santana, P.M. Forget, C. A. Kita, N. Lotfi, I. C. S. Machado, R. L. Muylaert, F. A. Rodrigues, T. M. Straka, and C. F. Dormann. 2025. A novel semantic theory of the assembly rules of interaction networks. EcoEvoRxiv. 10.32942/X20Q0V.

Morandin, L. A., and C. Kremen. 2013. Hedgerow restoration promotes pollinator populations and exports native bees to adjacent fields. Ecological Applications 23:829–839.

Nagano, Y., T. Yokoi, H. Taki, and T. Miyashita. 2025. Set-aside of grassland field margins enhances buckwheat pollination services in small-holder agricultural landscapes. Agriculture, Ecosystems & Environment 387:109628.

Nelder, J. A., and R. W. M. Wedderburn. 1972. Generalized Linear Models. Journal of the Royal Statistical Society. Series A (General) 135:370.

Peng, S., A. M. Ellison, and C. C. Davis. 2025. Climate change intensifies plant–pollinator mismatch and increases secondary extinction risk for plants in northern latitudes. Proceedings of the National Academy of Sciences of the United States of America 122:e2506265122.

Potts, S. G., J. C. Biesmeijer, C. Kremen, P. Neumann, O. Schweiger, and W. E. Kunin. 2010. Global pollinator declines: trends, impacts and drivers. Trends in Ecology & Evolution 25:345–353.

Potts, S. G., V. Imperatriz-Fonseca, H. T. Ngo, M. A. Aizen, J. C. Biesmeijer, T. D. Breeze, L. V. Dicks, L. A. Garibaldi, R. Hill, J. Settele, and A. J. Vanbergen. 2016. Safeguarding pollinators and their values to human well-being. Nature 540:220–229.

Pywell, R. F., M. S. Heard, B. A. Woodcock, S. Hinsley, L. Ridding, M. Nowakowski, and J. M. Bullock. 2015. Wildlife-friendly farming increases crop yield: Evidence for ecological intensification. Proceedings of the Royal Society B: Biological Sciences 282.

Rachersberger, M., G. D. Cordeiro, I. Schäffler, and S. Dötterl. 2019. Honeybee Pollinators Use Visual and Floral Scent Cues to Find Apple (Malus domestica) Flowers. Journal of Agricultural and Food Chemistry 67:13221–13227.

Requier, F., N. Pérez-Méndez, G. K. S. Andersson, E. Blareau, I. Merle, and L. A. Garibaldi. 2023. Bee and non-bee pollinator importance for local food security. Trends in Ecology & Evolution 38:196–205.

Rodríguez-Gasol, N., M. Viketoft, E. Chapurlat, J. A. Stenberg, M. Jonsson, and O. Lundin. 2026. Perennial flower strips increase pollinator and natural enemy abundance but have limited effects on pest control in adjacent crops. Agriculture, Ecosystems & Environment 396:110025.

Servedio, M. R., Y. Brandvain, S. Dhole, C. L. Fitzpatrick, E. E. Goldberg, C. A. Stern, J. Van Cleve, and D. J. Yeh. 2014. Not Just a Theory—The Utility of Mathematical Models in Evolutionary Biology. PLOS Biology 12:e1002017.

Shoemaker, L. G., J. A. Walter, L. A. Gherardi, M. H. DeSiervo, and N. I. Wisnoski. 2021. Writing mathematical ecology: A guide for authors and readers. Ecosphere 12:e03701.

Siopa, C., L. G. Carvalheiro, H. Castro, J. Loureiro, and S. Castro. 2024. Animal-pollinated crops and cultivars—A quantitative assessment of pollinator dependence values and evaluation of methodological approaches. Journal of Applied Ecology 61:1279–1288.

Sommer, J., V. Rao, and J. Sprayberry. 2022. Deconstructing and contextualizing foraging behavior in bumble bees and other central place foragers. Apidologie 53:32.

Tong, Z. Y., L. Y. Wu, H. H. Feng, M. Zhang, W. S. Armbruster, S. S. Renner, and S. Q. Huang. 2023. New calculations indicate that 90% of flowering plant species are animal-pollinated. National Science Review 10:219.

Turo, K. J., J. R. Reilly, T. P. M. Fijen, A. Magrach, and R. Winfree. 2024. Insufficient pollinator visitation often limits yield in crop systems worldwide. Nature Ecology and Evolution 8:1612–1622.

United Nations. 2015. Transforming our world: The 2030 Agenda for Sustainable Development.

